# GutCyc: a Multi-Study Collection of Human Gut Microbiome Metabolic Models

**DOI:** 10.1101/055574

**Authors:** Aria S. Hahn, Tomer Altman, Kishori M. Konwar, Niels W. Hanson, Dongjae Kim, David A. Relman, David L. Dill, Steven J. Hallam

**Affiliations:** Department of Microbiology and Immunology, University of British Columbia, Vancouver, BC. Canada; Koonkie Inc., Menlo Park, CA, USA; Biomedical Informatics, Stanford University School of Medicine, Stanford, CA 94305, USA; Whole Biome, Inc., 953 Indiana Street, San Francisco, CA 94107, USA; Computer Science and Artificial Intelligence Laboratory, Massachusetts Institute of Technology, Cambridge, MA 02139, USA; Department of Computer Science, University of British Columbia, Vancouver, BC, Canada; Department of Microbiology and Immunology, 299 Campus Drive, Stanford University School of Medicine, Stanford, CA 94305, USA; Department of Medicine, Stanford University School of Medicine, Stanford, CA 94305, USA; Veterans Affairs Palo Alto Health Care System, Palo Alto, CA 94304, USA; Department of Computer Science, Stanford University, Stanford, CA 94305, USA; Ecosystem Services, Commercialization and Entrepreneurship (ECOSCOPE), University of British Columbia, Vancouver, BC. Canada

## Abstract

Advances in high-throughput sequencing are reshaping how we perceive microbial communities inhabiting the human body, with implications for therapeutic interventions. Several large-scale datasets derived from hundreds of human microbiome samples sourced from multiple studies are now publicly available. However, idiosyncratic data processing methods between studies introduce systematic differences that confound comparative analyses. To overcome these challenges, we developed GUTCYC, a compendium of environmental pathway genome databases constructed from 418 assembled human microbiome datasets using METAPATHWAYS, enabling reproducible functional metagenomic annotation. We also generated metabolic network reconstructions for each metagenome using the PATHWAY TOOLS software, empowering researchers and clinicians interested in visualizing and interpreting metabolic pathways encoded by the human gut microbiome. For the first time, GUTCYC provides consistent annotations and metabolic pathway predictions, making possible comparative community analyses between health and disease states in inflammatory bowel disease, Crohn’s disease, and type 2 diabetes. GUTCYC data products are searchable online, or may be downloaded and explored locally using METAPATHWAYS and PATHWAY TOOLS.

## Background & Summary

The myriad collections of microorganisms found on and in the human body are known as the human microbiome [60]. Changes in microbiome structure and function have been implicated in numerous disease states including inflammatory bowel disease, cancer, and even cardiovascular disease [34, 11]. Increasingly, researchers are using high-throughput sequencing approaches to study the genes and genomes of microbiomes and characterize diversity and metabolic potential in relation to health and disease states [69] opening new opportunities for prevention and therapeutic intervention at the interface of microbial ecology, bioinformatics and medicine. The most densely colonized human habitat is the distal gut, inhabited by thousands of diverse microorganisms, as differentiated at the strain level. Despite providing essential ecosystem services, including nutritional provisioning, detoxification and immunological conditioning, the metabolic network driving matter and energy transformations by the distal gut microbiome remains largely unknown. Several large-scale metagenomic datasets (derived from hundreds of microbiome samples) from the Human Microbiome Project (HMP) [55], Beijing Genomics Institute (BGI) [58], and Metagenomes of the Human Intestinal Tract project (MetaHIT) [57] are now available on-line, creating an opportunity for large-scale metabolic network comparisons.

While the studies cited above provide the sequencing data, they do not provide the software environment used for generating their annotations. In contrast to these proprietary pipelines, over the past few years a number of metagenomic annotation pipelines available to third parties have emerged including IMG/M [46], Metagenome Rapid Annotation using Subsystem Technology (MG-RAST) [68], SMASHCOMMUNITY [9] and HUMANN [6]. Differing pipelines used to process sequence information between studies introduces biases based on idiosyncratic formatting, and alternative annotations or algorithmic methods. Specifically, support for metabolic pathway annotation varies significantly among pipelines due to differences in reference database selection with resulting impact on metabolic network comparisons. The most common metabolism reference database currently in use is Kyoto Encyclopedia of Genes and Genomes (KEGG) [26]. Although extant pipelines often provide links to KEGG module and pathways maps [26] (using KEGG Orthology (KO) or pathway identifiers) that can be visualized with coverage or gene count information using programs like KEGG Atlas [50], they do so using often incompatible formats. Such mapping is limited because there is no simple way to query, manipulate, or visualize the underlying implicit metabolic model directly. Moreover, prediction using KEGG results in amalgamated pathways with limited taxonomic resolution, impeding enrichment and association studies [6].

In responding to the deficiencies of existing tools, we recently developed a modular annotation and analysis pipeline enabling reproducible research [12] called METAPATHWAYS, that guides construction of Environmental Pathway/Genome Database (ePGDB)s from environmental sequence information [37] using PATHWAY TOOLS [27] and METACYC [32, 13, 14]. PATHWAY TOOLS is a production-quality software environment developed at SRI that supports metabolic inference and flux balance analysis based on the METACYC database of metabolic pathways and enzymes representing all domains of life. Unlike KEGG, METACYC emphasizes smaller, evolutionarily conserved or co-regulated units of metabolism and contains the largest collection (over 2,400) of experimentally validated metabolic pathways [7]. Navigable and extensively commented pathway descriptions, literature citations, and enzyme properties combined within an ePGDB provide a coherent structure for exploring and interpreting predicted metabolic networks from the human microbiome across multiple levels of biological information (DNA, RNA, protein and metabolites). Over 9,800 Pathway/Genome Database (PGDB)s have been developed by researchers around the world, and thus ePGDBs represent a data format for metabolic reconstructions that exhibit a potential for reusability in further studies.

Here we present GUTCYC, a compendium of over 418 ePGDBs constructed from public shotgun metagenome datasets generated by the HMP [55], the MetaHIT inflammatory bowel disease study [57], and the BGI diabetes study [58]. Relevant pipeline modules are summarized in Figure 1. GUTCYC provides consistent taxonomic and functional annotations, facilitates large-scale and reproducible comparisons between ePGDBs, and directly links into robust software and database resources for exploring and interpreting metabolic networks. This metabolic network reconstruction provides a multidimensional view of the microbiome that invites discovery and collaboration [30].

**Figure 1:**
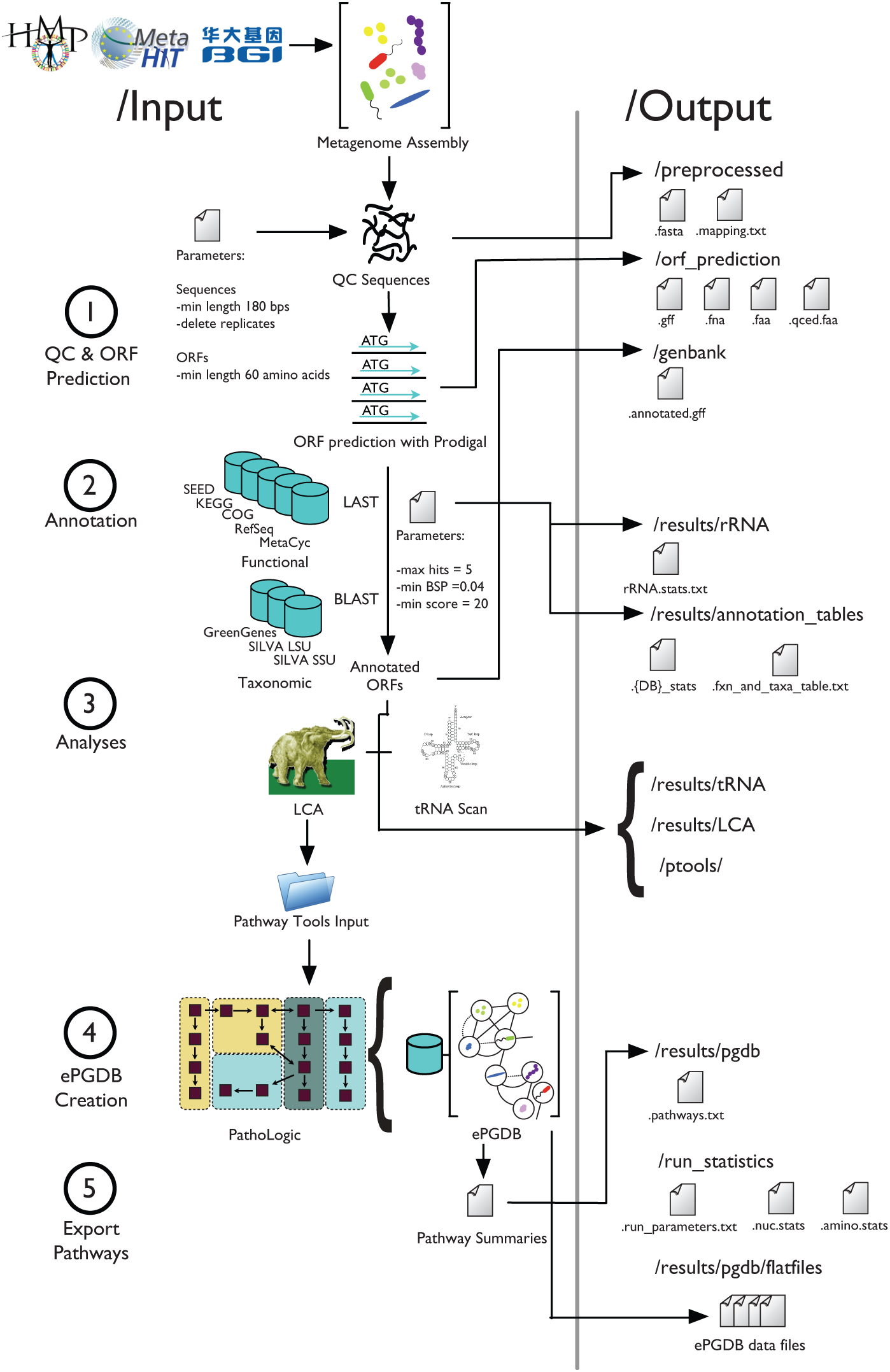
The METAPATHWAYS pipeline consists of five modular stages including (1) Quality control (QC) and ORF prediction (2) Functional and taxonomic annotation, (3) Analysis (4) ePGDB construction, and (5) Pathway export. Inputs and programs are depicted on the left with corresponding output directories and exported files on the right.

## Methods

### Metagenomic Data Sources

We collected 418 assembled human gut shotgun metagenomes from public repositories and supplementary materials sourced from the HMP (American healthy subjects, *n* = 148) [55], a MetaHIT (European inflammatory bowel disease subjects, *n* = 125) [17], and a BGI (Chinese type 2 diabetes, n = 145) study [58]. See Supplementary Table 1 for a detailed listing of accession numbers and file descriptors.

### Data Processing

Microbiome project sample metadata were manually curated to ensure compatibility with METAPATHWAYS. ePGDBs were created for each sample by running the METAPATHWAYS 2.5 pipeline and the PATHWAY TOOLS version 17.5, using the assembled metagenomes described above. The pipeline consists of five modular steps, including (1) quality control and ORF prediction, (2) homology-based functional and taxonomic annotation, (3) analyses consisting of tRNA and lowest common ancestor (LCA) [24] identification, (4) construction of ePGDBs using PATHWAY TOOLS and, finally, (5) pathway export [38, 33] (see Figure 1). The following paragraphs describe the individual processing steps required to construct an ePGDB for each sample, starting with assembled contigs in FASTA format.

#### Quality Control

Contigs from each sample were collected from their respective repositories and curated locally. The MetaPathways pipeline performs a number of quality control steps. First, each contig was checked for the presence of ambiguous base pairs and homopolymer runs, splitting contigs into smaller sequences by removing such problematic regions. Next, the contigs were screened for duplicates. Finally, a length cutoff of 180 base pairs was applied to the remaining sequences to ensure that very short sequences were removed from downstream processing steps [39].

#### ORF Prediction

Sequences passing quality control were scanned for ORFs using METAPRODIGAL [25], a robust ORF prediction tool for microbial metagenomes considered to be among the most accurate ORF predictors [65]. Resulting ORF sequences were translated to amino acid sequences using NCBI genetic code table 11 for bacteria, archaea, and plant plastids [8]. Translated amino acid sequences shorter than 30 amino acids were removed as these sequences approached the so-called functional homology search “twilight zone”, where it becomes difficult to detect true homologs [61].

#### Functional Annotation

The quality controlled amino acid sequences were queried against a panel of functionally-annotated protein reference databases including KEGG [26] (downloaded 2011-06-18), COG [63] (downloaded 2013-12-27), METACYC [14] (downloaded 2011-07-03), REFSEQ [62] (downloaded 201401-18), and SEED [52] (downloaded 2014-01-30). Protein sequence similarity searches were performed using the program FAST [42] with standard alignment result cutoffs (E-value less than 1 × 10^−5^, bit-score greater than 20, and alignment length greater than 40 amino acids [61]; and BLAST-score ratio (BSR) greater than 0.4 [59]). The choice of parameter thresholds were selected to maximize annotation accuracy, and were guided based on parameter choices used in previous studies [22, 70, 67].

#### Taxonomic Annotation

Quality-controlled contigs were also searched against the SILVA [56] (version 115) and GREENGENE [16] (downloaded 2012-11-06) ribosomal RNA (rRNA) gene databases using BLAST version 2.2.25, with the same post-alignment thresholds applied as was previously described for the functional annotation. BLAST was applied for 16S annotation because it has greater sensitivity for nucleotide-nucleotide searches than FAST, and the smaller reference database sizes make the relatively larger computational requirement justifiable.

Additionally, predicted ORFs were taxonomically annotated using the LCA algorithm [24] for taxonomic binning. In brief, the LCA in the NCBI Taxonomy Database (TaxonDB) [62] was selected based on the previously calculated FAST hits from the REFSEQ database. This effectively sums the number of FAST hits at the lowest shared position of the TaxonDB. The RefSeq taxonomic names often contain multiple synonyms or alternative spellings. Therefore, names that conform to the TaxonDB were selected in preference over unknown synonyms.

#### tRNA Scan

MetaPathways uses TRNASCAN-SE version 1.4 [44] to identify relevant tRNAs from quality-controlled sequences. Resulting tRNA identifications are appended as additional functional annotations.

#### ePGDB Creation

Functional annotations were parsed and separated into three files that serve as inputs to PATHWAY TOOLS, namely: (1) an annotation file containing gene product information (**`0.pf`**), (2) a catalog of contigs and scaffolds (**`genetic-elements.dat`**), and (3) a PGDB parameters file (**`organism-params.dat`**). The PathoLogic module [19, 15] in the PATHWAY TOOLS software, was used to build the ePGDB and predict the presence of metabolic pathways based on functional annotations. Following ePGDB construction, the base pathways (i.e., pathways that are not contained within superpathways) were extracted from ePGDBs to generate a summary table of predicted metabolic pathways for each sample.

#### Accessibility and Flexibility

METAPATHWAYS 2.5 generates data in a consistent file and directory structure. The output for each sample is contained within a single directory, which in turn is organized into sub-directories containing relevant files (see Figure 1). The METAPATHWAYS 2.5 graphical user interface (GUI) enables interactive exploration of individual sample results along with comparative queries of multiple samples, and is designed for fast and interactive data visualization and searches *via* a custom knowledge engine data structure. Input and output files are available for download from the GUTCYC website (**www.gutcyc.org**) and may be readily explored in the METAPATHWAYS GUI or PATHWAY TOOLS on LINUX, MAC OS X and WINDOWS machines.

#### Computational Environment

Computational processing was performed using a local cluster of machines in the Hallam laboratory and on the BUGABOO cluster on the Canadian WestGrid computation resource [5]. The Hallam lab computers have a configuration profile of 2×2.4 GHz Quad-Core Intel Xeon processors with 64 GB 1066 MHz DDR3 RAM. The BUGABOO cluster provides 4,584 cores with 2 GB of RAM per core on average. The average sample took 7-8 hours to process on a single thread, and the span of the processing required to generate the GUTCYC COLLECTION was 135 days.

### Software Availability

METAPATHWAYS 2.5, including integrated third party software, is available on GitHub, including both software [2] (licensed under the GNU General Public License, version 3), and a tutorial [3] released under the Creative Commons Attribution License (allows reuse, distribution, and reproduction given proper citation). PATHWAY TOOLS is available under a free license for academic use, and may be commercially licensed [4]. METAPATHWAYS outputs were processed using PATHWAY TOOLS version 17.5 under default settings except for disabling of the PathoLogic taxonomic pruning step (i.e., **`-no-taxonomic-pruning`**) as was described previously [22], and an additional refinement step of running the Transport Inference Parser [43] to predict transport reactions (i.e., **`-tip`**). FAST is freely available under a (licensed under the GNU General Public License, version 3) software license on our GitHub page [1].

### Data Records

A list of each sample, provenance, and relevant data processing steps can be found in Supplementary Table 1. All records are available at the GUTCYC project website (**www.gutcyc.org**). Each sample’s data records are contained within a single directory. Within this directory, sub-directories and files are located as depicted in Figure 1. A summary of the data present in the GUTCYC COLLECTION is presented in Table 1. A full set of summary data for each ePGDB may be found in Supplementary Table 2.

**Table 1:** Summary statistics for the GUTCYC COLLECTION across 418 samples. The statistics for the number of bases processed is in units of Megabases. “Func. Annots.”: functional annotations. “Trans. Reactions” are transport reactions. “Compounds” are small molecule metabolites. “Base Pathways” include all pathways except complex pathways known as Super-Pathways.

|  | Min | 1<sup>st</sup> Quartile | Median | 3<sup>rd</sup> Quartile | Max |
| --- | --- | --- | --- | --- | --- |
| Bases | 0.98 | 54.75 | 81.35 | 113.75 | 370.51 |
| Contigs | 2,506 | 27,788 | 47,486.5 | 76,275.75 | 399,331 |
| ORFs | 2,448 | 61,703.5 | 95.531 | 139,690 | 550,312 |
| Func. Annots. | 2,176 | 57,102.25 | 86,054.5 | 123,747.25 | 425,033 |
| Reactions | 1,635 | 2,385.5 | 3,438 | 3,667.75 | 4,881 |
| Trans. Reactions | 12 | 26 | 31 | 34 | 46 |
| Compounds | 1,052 | 1,678 | 2,008.5 | 2,119.5 | 2,676 |
| Base Pathways | 257 | 350 | 616 | 654 | 832 |

#### preprocessed

For a sample with an identifier of **`<sample_ID>`**, this directory contains two files: (1) **`<sample_ID>.fasta`**, which contains the renamed, quality-controlled sequences, and (2) **`<sample_ID>.mapping.txt`**, which maps the original sequence names to the new names assigned by METAPATHWAYS.

Sequences are renamed to **`<sample_ID>_X`** where *X* is the zero-indexed contig number pertaining to the order in which the contig appears in the input file (e.g., a contig identified as DLF001_27 is interpreted as the 28^th^ contig listed in the FASTA file for sample DLF001’s assembly).

#### orf_prediction

This directory contains four files, (1) **`<sample_ID>.fna`** which contains nucleic acid sequences of all predicted ORFs, (2) **`<sample_ID>.faa`** which contains amino acid sequences of all predicted ORFs, (3) **`<sample_ID>.qced.faa`** which contains amino acid sequences of all predicted ORFs meeting user defined quality thresholds (in this study, a minimum length of 60 amino acids), and (4) **`<sample_ID>.gff`**, a General Feature Format (GFF) file [18] containing all quality-controlled sequences and information about the strand (− or +) on which the ORF was predicted. ORFs are named **`<sample_ID>_X_Y`**, where X is the contig number pertaining to the order in which the contig appears and *Y* represents the order in which the ORFs were predicted on the contig.

#### results

This directory contains four sub-directories: (1) **`annotation_table`**, (2) **`rRNA`**, (3) **`tRNA`**, and (4) **`pgdb`**. The **`annotation_table`** sub-directory contains (1) statistics files for each functional database used to annotate the ORFs (**`<sample_ID>.<DB>_stats_<index>.txt`**), (2) **`<sample_ID>.functional_and_taxonomic_table.txt`** detailing the length, location, strand and annotation (functional and taxonomic) of each ORF, and (3) a file listing all ORFs and their functional annotations (**`<sample_ID>.ORF_annotation_table.txt`**). The prokaryotic 16S ribosomal RNA gene is a standard marker gene used for measuring taxonomic diversity [66]. The **`rRNA`** sub-directory contains files detailing statistics for each taxonomic database used to annotate the ORFs (named as **`<sample_ID>.<DB>.rRNA.stats.txt`**). The **`tRNA`** sub-directory contains (1) **`<sample_ID>.trna.stats.txt`**, detailing the type, anticodon, location and strand of each predicted tRNA and (2) **`<sample_ID>.tRNA.fasta`** containing all predicted tRNA sequences. The **`pgdb`** sub-directory contains a **`<sample_ID>.pwy.txt`** file describing metabolic pathways predicted in the ePGDB, specifically, each predicted pathway, the ORF identities involved in each pathway, the enzyme abundance, and the pathway coverage in a tabular format navigable via the METAPATHWAYS GUI.

#### genbank

This directory contains a file named **`<sample_ID>.annotated.gff`**, a GFF file containing all quality-controlled sequences with their annotations.

#### ptools

This directory contains the three files necessary to build a ePGDB using PATHWAY TOOLS: (1) **`genetic-elements.dat`**, (2) **`organism-params.dat`**, and (3) **`0.pf`** which contains all functional annotations to be processed by PATHWAY TOOLS. A sub-directory called **`flat-files`** contains flat files describing database objects such as compounds, reactions, pathways (each of which is described in more detail in [28]) for individual ePGDBs.

#### run_statistics

This directory contains three files: (1) **`<sample_ID>.run.stats`**, the parameters used to process the sample; (2) **`<sample_ID>.nuc.stats`**, the number and length of nucleic acid sequences before and after quality control filtering; and (3) **`<sample_ID>.amino.stats`**, the number and length of amino acid sequences before and after quality control filtering.

## Technical Validation

GUTCYC was derived from third-party sequence data from three publicly-available human gut microbiome sampling projects with metagenomic assemblies, with published details on their own technical validation steps: the HMP [55], a MetaHIT study [17], and a BGI study [58]. The technical validation of third-party software used in METAPATHWAYS may be found in the corresponding publications for METAPRODIGAL [25], BLAST [10], and `T`RNASCAN-SE [44]. GUTCYC functional sequence similarity was computed using FAST, an aligner based on a version of LAST [35], with multi-threading performance improvements and new support for generating BLAST-like E-values, with significant correlation with BLAST output (*R*^2^ = 0.887, *P* < 0.01) [38]. Validation of the overall METAPATHWAYS pipeline may be found in previously published reports [22, 23] with specific emphasis on how changes in taxonomic pruning, read length and metagenomic assembly coverage impact the accuracy and sensitivity of pathway recovery. In brief, pathway prediction is affected by taxonomic distance, sequence coverage and sample diversity, nearing an asymptote of maximum accuracy for metagenomes with increasing coverage. Additionally, like any alignment-based analysis, annotation quality is a function of both the level of errors in the input sequence data and the selection of reference databases. Summary data generated for each ePGDB as presented in Supplementary Table 2 was reviewed to detect samples with unusual statistics, such as a lack of 16S gene annotations. The metabolic reconstruction pathways were computationally predicted using the Pathway Tools PathoLogic module [53], which has an accuracy of 91% [15]).

## Usage Notes

Once a set of data such as GUTCYC COLLECTION has been crafted into a format that is both comprehensible to domain experts, and interpretable by machines, there are myriads of uses that can be explored. For example, comparing ePGDBs with sets of microbial PGDBs from the same environment can aid in identifying “distributed pathways” present in the metagenome metabolic reconstruction, but absent from any individual genomic metabolic reconstruction [22]. The predicted transport proteins can be used to predict trophism patterns within a community. Furthermore, the PATHWAY TOOLS software allows for sophisticated comparative analyses between ePGDBs, at the level of compounds, reactions, enzymes, and pathways [31]. The METAFLUX [40] module of PATHWAY TOOLS for performing flux balance analysis (FBA) [51] can be used with GUTCYC ePGDBs to generate quantitative simulations of microbiome growth and pathway flux. A set of microbiome metabolic models also facilitates the exploration of the impact of xenobiotics [21], and provides a computational substrate for systems biology approaches to engineering the gut microbiome [47]. Figure 2 demonstrates the user interface for METAPATHWAYS and PATHWAY TOOLS, along with example data analysis use cases.

**Figure 2:**
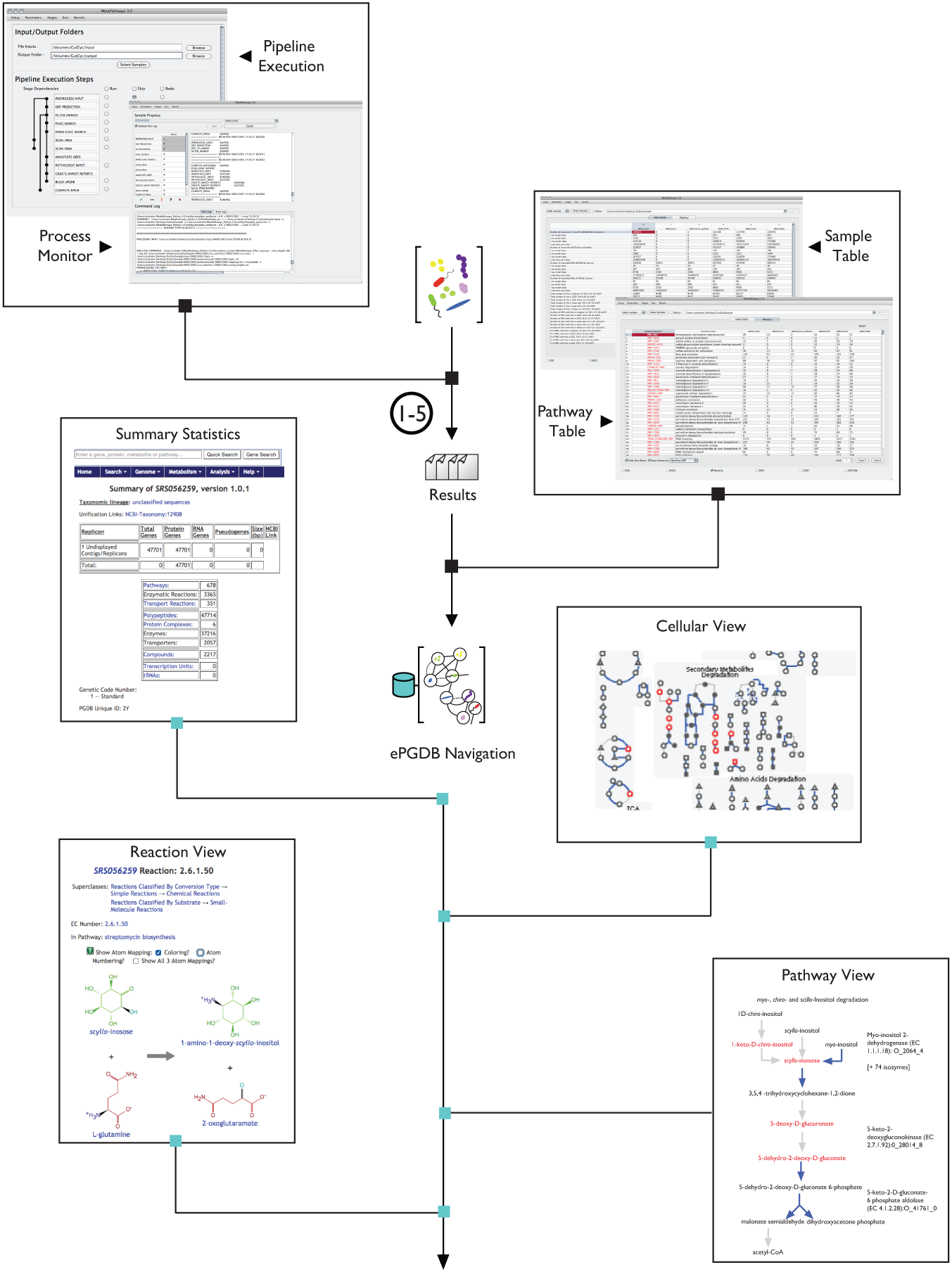
GUTCYC ePGDB use cases. In the upper left and upper right insets, a GUTCYC ePGDB is opened in METAPATHWAYS. In the upper left, we display the Pipeline execution step, and the Process Monitor interfaces. In the upper right, we display the Summary Table (with exportable sample statistics), and the Pathway Table (with exportable pathway abundances) interfaces. In the lower for inset images, a GUTCYC ePGDB is opened in PATHWAY TOOLS. Clockwise from the upper left, we display the ePGDB summary statistics, interative metabolic network visualization, the Pathway View, and the biochemical Reaction View.

In this section we motivate further two specific use cases for GUTCYC. In the first case, we demonstrate how to use a GUTCYC ePGDB to determine the metabolic path between two small molecules of interest. In the second case, we use GUTCYC to visualize different levels of biological information, e.g. metabolomics data, in the context of a microbiome metabolic network.

### Optimal Metabolite Tracing

The PATHWAY TOOLS software provides advanced biochemical querying capabilities for both PGDBs and ePGDBs. One such capability is energy-optimal metabolite tracing. To summarize, given both a starting and a terminal/target compound within an ePGDB, PATHWAY TOOLS is able to compute the shortest and most energetically-favorable route through the metabolic reaction network. While there is no guarantee that, in a complex milieu such as the gut microbiome, the syntrophic flux will necessarily follow a short and minimal energy path, these criteria allow us to narrow down a multiplicity of possible paths to a single parsimonious candidate path.

In a study by Koeth et al., they demonstrated a causal connection between the intestinal gut microbiota’s metabolism of red meat and the promotion of atherosclerosis [36]. In brief, the gut microbiome is capable of transforming excess *L*-carnitine into trimethylamine (TMA), which is further processed by the liver to form the cardiovascular disease-associated metabolite trimethylamine *N*-oxide (TMAO). Using this biotransformation as a motivating case, we queried the GUTCYC SRS015217CYC ePGDB for the biochemical reaction path from *L*-carnitine to TMA, which is not provided explicitly by Koeth et al. Utilizing the PATHWAY TOOLS Metabolic Route Search feature, we found an optimal path between L-carnitine to TMA, using the METACYC *carnitine degradation II* pathway (PWY-3602, expected in *Proteobacteria*) along with a betaine reductase reaction (EC 1.21.4.4; found in *Clostridium sticklandii* and *Eubacterium acidaminophilum*, both species affiliated with the order Clostridiales), minimizing the number of enzymes involved and chemical bond rearrangements. PATHWAY TOOLS found the optimal path in seconds, displayed in Figure 2.

*L*-carnitine and glycine betaine have known transporter families that facilitate their movement across the cell membrane [48], as do TMA and TMAO [49], and thus this metabolic route may be a distributed pathway [22]. In fact, no single PGDB in the BIOCYC COLLECTION of over 5,500 microbial genomes (release 19.0 [14]), has both the *carnitine degradation II* pathway and the *betaine reductase* reaction, which suggests that there is no single microbe capable of completing this entire metabolic route.

The metabolic route identified may also help generate mechanistic hypotheses from microbiome study observations. Among the findings reported in [36] in the Supplementary Materials is that *all statistically-significant correlations (positive or negative) found between plasma TMAO levels and species abundance, involved species affiliated wth the order Clostridiales,* which is the subsuming taxon of the betaine reductase reaction’s taxonomic range, as curated in METACYC. This indicates that Clostridiales are integral to understanding the regulation of TMA and TMAO concentrations in the gut, which in turn affects plasma concentrations. This demonstrates the power of ePGDBs in computing connections between nutritional or pharmaceutical inputs (such as *L*-carnitine) to identify potential interactions with known disease biomarkers (as TMAO is to cardiovascular disease).

### High-Throughput Data Visualization

Another capability of PATHWAY TOOLS is to visualize the results of high-throughput experiments mapped onto the Cellular, Genome, and Regulation Overviews, or as “Omics Pop-Ups” when viewing a particular pathway [54]. Specifically, PATHWAY TOOLS provides support for the analysis of mass spectrometry data, by automatically mapping a list of monoisotopic masses to matching entries in METACYC, or in specific ePGDBs [29]. As a demonstration of this capability, we analyzed mass-spectrometry data from a metabolomic study of humanized mice microbiomes [45]. The dataset contained 867 unique masses, of which 453 masses were identified using METACycby performing standard adduct monoisotopic mass manipulations [64], followed by monoisotopic mass search using Pathway Tools. We mapped the identified compounds on the GUTCYC Cellular Overview [41], as seen in Figure 3. This facilitates turning a massive table of data into a more intuitive construct based on the community metabolic interaction network, enabling more efficient pattern matching. For example, using the enrichment analysis tools in Pathway Tools [29], we identified the pathway class of “Secondary Metabolites Degradation” as enriched for identified compounds (*p* = 2.0 × 10^−2^, Fisher Exact Test with Benjamini-Hochberg multiple testing correction). By visually inspecting the pathways in the class, we can see that pathway P562-PWY, “myo-, chiro-, and scillo-inositol degradation pathway”, has four matched compounds from the metabolomics dataset.

**Figure 3:**
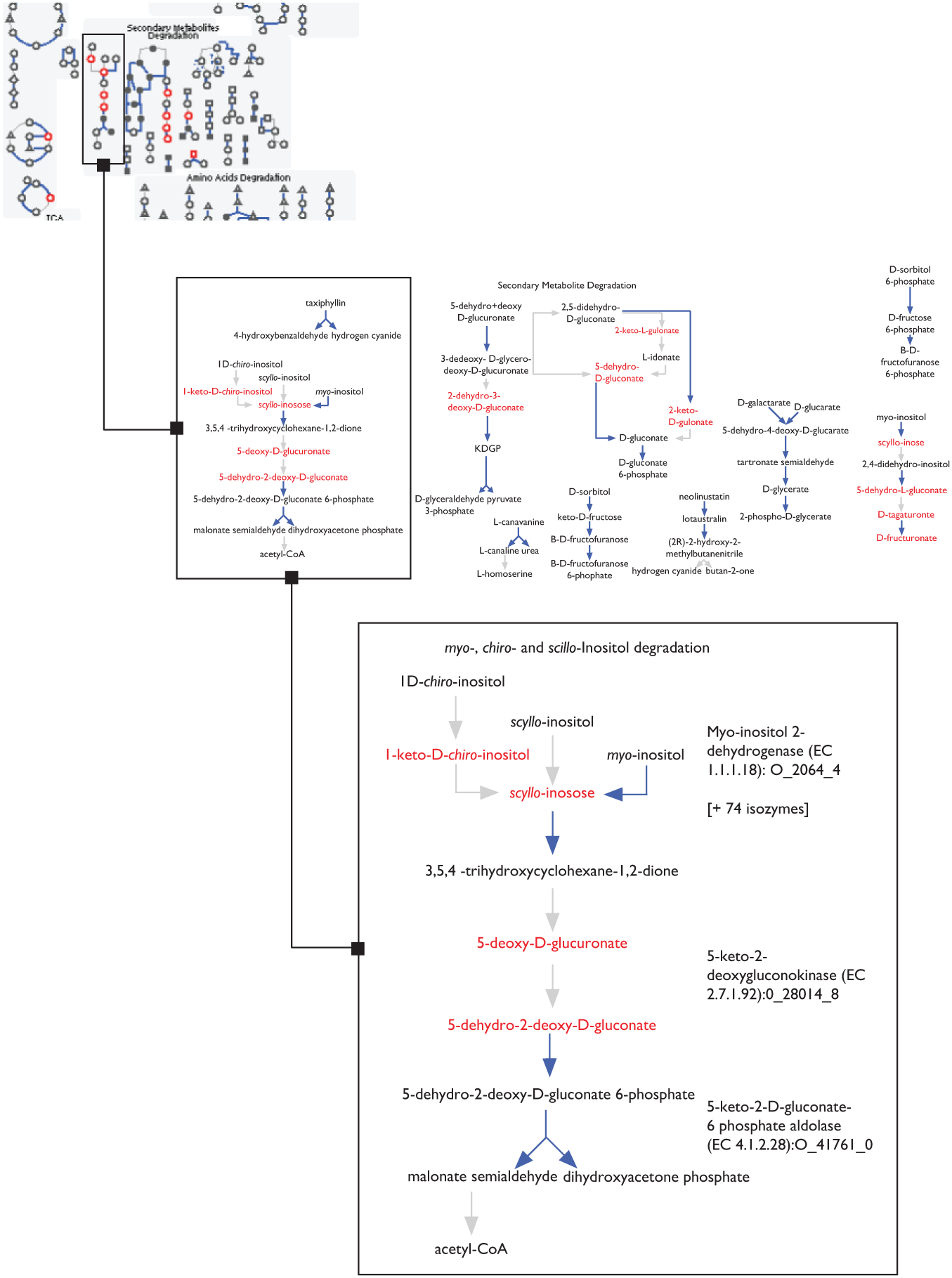
The Cellular Overview for the SRS056259CYC ePGDB, at three different zoom levels, with compounds highlighted in red if identified from a mass spectrometry analysis of the gut microbiome [45]. Compounds with no mass spectrometry highlights appear as grey icons. Reactions with enzyme data in SRS056259CYC are drawn in blue. The top left inset shows a fraction of the full metabolic map. The middle inset shows a zoom-in of the “Secondary Metabolite Degradation” pathway class. Bottom right inset shows zoom-in on Pathway P562-PWY, “myo-, chiro-, and scillo-inositol degradation pathway”, showing four mass-spectrometry identified compounds in red.

## Acknowledgments

We would like to thank Peter D. Karp for feedback on the MetaPathways software and the GutCyc project; Robert Pesich for orchestrating our sneakernet transfer of data; and Les Dethlefsen for assisting in loading the data onto the Relman Lab server. A special thanks to the members of the Hallam, Relman, and Dill labs, and Whole Biome, for constructive feedback on the GUTCYC project. Thank you to Pallavi Subhraveti of SRI International for help with exporting GUTCYC data using Pathway Tools. Thank you to the Stanford FarmShare computation resource, for aiding in the development of an early version of GUTCYC.

The GutCyc project at UBC was carried out under the auspices of Compute/Calcul Canada, Genome Canada, Genome British Columbia, Genome Alberta, the Natural Science and Engineering Research Council (NSERC) of Canada, Ecosystem Services, Commercialization Platforms and Entrepreneurship (ECOSCOPE) program, the Canadian Foundation for Innovation (CFI), and the Canadian Institute for Advanced Research (CIFAR) through grants awarded to SJH. ASH was supported by the Alexander Graham Bell Canada Graduate Scholarships-Doctoral Program (CGS D) administered by NSERC. KMK was supported by the Tula Foundation funded Centre for Microbial Diversity and Evolution (CMDE) at UBC. NWH was supported by a four year doctoral fellowship (4YF) administered through the UBC Faculty of Graduate and Postdoctoral Studies. TA was partially supported by the Stanford University School of Medicine Dean’s Funds and the NIH Biotechnology Training Grant at Stanford (grant number 5T32 GM008412). TA and DLD were partially supported by a King Abdullah University of Science and Technology (KAUST) research grant under the KAUST Stanford Academic Excellence Alliance program. DAR was supported by NIH/NIGMS 5R01GM099534 and by the Thomas C. and Joan M. Merigan Endowment at Stanford University. Additional computational resources were provided gratis through the Stanford FarmShare resource.

## Author Contributions

Tomer Altman and Steven Hallam conceived of the GutCyc project as part of a movement to develop the Environmental Genome Encyclopedia (EngCyc): a compendium of microbial community metabolic blueprints supported by high performance software tools on grids and clouds. Niels Hanson, Kishori Konwar, Aria Hahn and Dongjae Kim developed the MetaPathways software pipeline with direction from Steven Hallam and assistance from Tomer Altman and others at SRI International. Aria Hahn, and Kishori Konwar compiled the microbiome sequence datasets, constructed GutCyc ePGDBs and created figures for the manuscript. Tomer Altman generated validation datasets and drafted an early version of the manuscript with Aria Hahn and Steven Hallam. Dongjae Kim developed the GutCyc website. All authors contributed to the final preparation of the manuscript. Steven Hallam, David L. Dill and David A. Relman supervised the project. All authors reviewed and approved the final manuscript.

## Data Availability

The GUTCYC COLLECTION, along with metadata for all samples, is freely available at **www.gutcyc.org**. See below for the figshare DOI Data Citation.

## Glossary

BGI: Beijing Genomics Institute. 2, 3, 9
BSR: BLAST-score ratio. 5
ePGDB: environmental Pathway/Genome Database. 3-13
FBA: flux balance analysis. 10
GFF: General Feature Format. 8
GUI: graphical user interface. 6, 8
HMP: Human Microbiome Project. 2, 3, 9
KAUST: King Abdullah University of Science and Technology. 14
KEGG: Kyoto Encyclopedia of Genes and Genomes. 2, 3
KO: KEGG Orthology. 2
LCA: lowest common ancestor. 5, 6
MetaHIT: Metagenomes of the Human Intestinal Tract project. 2, 3, 9
ORF: open reading frame. 3-6, 8
PGDB: Pathway/Genome Database. 3, 6, 9, 10, 13
TaxonDB: NCBI Taxonomy Database. 6
TMA: trimethylamine. 10, 13
TMAO: trimethylamine N-oxide. 10, 13

## Data Citations

Hahn, Aria S; Altman, Tomer; Konwar, Kishori M; Hanson, Niels W; Kim, Dongjae; Relman, David A; Dill, David L; Hallam, Steven J (2016): GutCyc. figshare. https://dx.doi.org/10.6084/m9.figshare.c.3283562. Retrieved: 16:04, Jul 12, 2016 (GMT).

